# Elimination of galanin synthesis in noradrenergic neurons reduces galanin in select brain areas and promotes active coping behaviors

**DOI:** 10.1101/651943

**Authors:** Rachel P. Tillage, Natale R. Sciolino, Nicholas W. Plummer, Daniel Lustberg, L. Cameron Liles, Madeline Hsiang, Jeanne M. Powell, Kathleen G. Smith, Patricia Jensen, David Weinshenker

**Author notes:** These authors contributed equally to the work (Rachel P. Tillage, Natale R. Sciolino). Co-corresponding authors: David Weinshenker, Ph.D., Department of Human Genetics, Emory University School of Medicine, Whitehead 301, 615 Michael St., Atlanta, GA 30322, and Patricia Jensen, Ph.D., Neurobiology Laboratory, National Institute of Environmental Health Sciences, National Institute of Health, Department of Health and Human Services, Research Triangle Park, NC 27709.

## Abstract

Accumulating evidence indicates that disruption of galanin signaling is associated with neuropsychiatric disease, but the precise functions of this neuropeptide remain largely unresolved due to lack of tools for experimentally disrupting its transmission in a cell type-specific manner. To examine the function of galanin in the noradrenergic system, we generated and crossed two novel knock-in mouse lines to create animals lacking galanin specifically in noradrenergic neurons (*Gal^cKO-Dbh^*). We observed reduced levels of galanin peptide in pons, hippocampus, and prefrontal cortex of *Gal^cKO-Dbh^* mice, indicating that noradrenergic neurons are a significant source of galanin to those brain regions, while midbrain and hypothalamic galanin levels were comparable to littermate controls. In these same brain regions, we observed no change in levels of norepinephrine or its major metabolite at baseline or after an acute stressor, suggesting that loss of galanin does not affect noradrenergic synthesis or turnover. *Gal^cKO-Dbh^* mice had normal performance in tests of depression, learning, and motor-related behavior, but had an altered response in some anxiety-related tasks. Specifically, *Gal^cKO-Dbh^* mice showed increased marble and shock probe burying and had a reduced latency to eat in a novel environment, indicative of a more proactive coping strategy. Together, these findings indicate that noradrenergic neurons provide a significant source of galanin to discrete brain areas, and noradrenergic-specific galanin opposes adaptive coping responses.

## INTRODUCTION

The neuropeptide galanin was first discovered in the early 1980s in porcine gut. Subsequent studies revealed that galanin is expressed throughout the brain and body of both humans and rodents (Tatemoto et al. 1983; Lang et al. 2015; Fang et al. 2015; Kofler et al. 2004) and modulates a variety of physiological processes including feeding, nociception, seizures, stress responses, cognition, and mood (Mitsukawa et al. 2008; Lang et al. 2015). Given this broad range of functions, it is not surprising that genome-wide association studies have implicated variants in genes encoding galanin and its receptors in conferring increased risk of depression and anxiety in humans (Wray et al. 2012; Juhasz et al. 2014; da Conceicao Machado et al. 2018). The exact role of galanin in these complex disorders, however, has yet to be determined.

Preclinical studies examining the effects of galanin have relied heavily on global knockouts or intracerebroventricular infusion of galanin agonists or antagonists. These experimental strategies lack regional specificity and often generate conflicting results. For example, galanin agonists administered into the rodent brain have been reported to elicit an anxiolytic effect in some tests (e.g. conflict test and elevated zero maze), and no effect in other anxiety assays (elevated plus maze, light-dark exploration, and open field) (Karlsson and Holmes 2006; Karlsson et al. 2005; Bing et al. 1993; Rajarao et al. 2007). Likewise, galanin agonists also produce mixed effects on behavior in the forced swim and tail suspension tests (Lu et al. 2005; Kuteeva et al. 2008a; Kuteeva et al. 2008b; Holmes et al. 2005; Bartfai et al. 2004). Furthermore, global galanin knockout mice exhibit a wide variety of phenotypes, including endocrine, neurological, and behavioral deficits (Holmes et al. 2000; Ahren et al. 2004; Wynick et al. 1998; Wynick and Bacon 2002; Zachariou et al. 2003; Adams et al. 2008). These findings suggest that galanin’s effects are specific to the cell type, brain region, circuit, and/or receptor(s) being engaged. Thus, strategies to manipulate galanin in a cell type-specific manner will help unravel its distinct functional roles in diverse physiological processes and disease states.

Galanin is expressed abundantly in a subset of noradrenergic neurons located in the locus coeruleus (LC) and subcoeruleus (SubC) in both rodents and humans (Holets et al. 1988; Skofitsch and Jacobowitz 1985; Perez et al. 2001; Le Maitre et al. 2013; Chan-Palay et al. 1990; Melander 1986). Intriguingly, transgenic mice overexpressing galanin in all noradrenergic neurons exhibit seizure resistance, mild cognitive deficits, and resilience against yohimbine-induced anxiety-like behavior (Steiner et al. 2001; Mazarati et al. 2000; Holmes et al. 2002). Because the transgene is expressed in all neurons that produce norepinephrine (NE) and epinephrine, and ectopically in some non-NE neurons (e.g. piriform cortex, entorhinal cortex) (Steiner et al. 2001), it is difficult to ascribe phenotypes to the subset of NE neurons that endogenously express galanin. The role of noradrenergic-derived galanin in normal and impaired brain function therefore remains unclear (for reviews see Weinshenker and Holmes 2015; Lang et al. 2015; Hökfelt et al. 2018; Sciolino and Holmes 2012).

To begin to address this important question, we combined two new mouse alleles - a conditional knockout allele of *Gal* and a knock-in cre driver allele under control of the noradrenergic-specific *Dbh* promoter - to generate a mouse model in which galanin is selectively disrupted in noradrenergic neurons. Using this model, we measured the proportion of noradrenergic neuron-derived galanin in discrete brain regions and determined the consequences of its loss on behaviors relevant to anxiety, depression, cognition, and gross motor function. Our biochemical findings reveal that noradrenergic neurons provide a significant source of galanin to the cortex and hippocampus, and that loss of galanin has no effect on NE levels or turnover. Furthermore, we demonstrate that noradrenergic neuron-specific galanin signaling plays a role in the regulation of defensive coping behaviors in the context of anxiogenic environmental manipulations.

## MATERIALS AND METHODS

### Animals

All procedures related to the use of animals were approved by the Animal Care and Use Committee of the NIEHS and Emory University and were in accordance with the National Institutes of Health guidelines for the care and use of laboratory animals. Mice were maintained on a 12/12 h light-dark cycle with access to food and water *ad libitum* unless stated otherwise. **Generation of mouse lines**

To generate the *Gal^cKO^* allele (*Gal^tm1a(KOMP)Pjen^*), we obtained a targeting vector (Project ID: CSD82928) containing loxP-flanked *Gal* exon 3 and FRT-flanked *LacZ* and *Neo* cassettes from the KOMP Repository (https://www.komp.org). Linearized vector was electroporated into G4 embryonic stem cell (George et al. 2007), and homologous recombinants were identified by Southern blotting with a *Neo* probe and long range PCR. Recombinant cells were injected into B6(Cg)-*Tyr^c-2J^*/J blastocysts to produce chimeric mice. Heterozygous offspring of the chimeras were crossed to B6.Cg-*Tg(ACTFlpe)9205Dym*/J mice (Rodriguez et al. 2000; Jackson Lab stock no. 005703) to excise the FRT-flanked *LacZ* and *Neo* cassettes and generate the conditional allele. The *Gal^cKO^* mice were thereafter maintained by backcrossing to C57BL/6J mice. *Cre*-mediated recombination of this allele deletes exon 3 and introduces a frameshift if exon 2 is spliced to exon 4, thus eliminating expression of both galanin and galanin message associated protein (GMAP), which are encoded by the same mRNA.

To generate the *Dbh^cre^* allele, we employed homologous recombination in G4 embryonic stem cells to insert a rox-flanked transcriptional stop cassette, *cre* cDNA, rabbit β-globin polyadenylation cassette, and attB/attP-flanked neomycin resistance cassette into the *Dbh* locus, immediately following the start codon. A recombinant clone was transiently transfected with pPGKPhiC31obpa (Raymond and Soriano 2007) to excise the neomycin resistance cassette before cells were injected into B6(Cg)-*Tyr^c-2J^* blastocysts to produce chimeric mice. Following germline transmission, we crossed heterozygotes with B6;129-*Tg(CAG-dre)1Afst* mice to permanently excise the rox-flanked stop cassette and generate the *Dbh^cre^* mouse line. The *Dbh^cre^* mice were thereafter maintained by backcrossing to C57BL/6J.

*Dbh^cre^; RC::LTG* mice were generated by crossing heterozygous *Dbh^cre^* to homozygous *RC::LTG* mice (Plummer et al. 2017)*. Gal^cKO-Dbh^* mutants were created by crossing *Dbh^cre^*^/+^;*Gal^cKO^*^/+^ double heterozygotes or *Dbh^cre^*^/+^;*Gal^cKO^* homozygotes to *Gal^cKO^* homozygotes (**Fig. 1a**). *Gal^cKO-Nestin^* mutants were created by crossing *Nestin^cre^*^/+^;*Gal^cKO^* homozygotes to *Gal^cKO^* homozygotes (*Nestin^cre^*^/+^ breeders were obtained from The Jackson Laboratory, Bar Harbor, ME, Stock No. 003771). *Gal^NULL^* mutants were generated by crossing *Gal^cKO^* heterozygotes to a FVB/N-Tg(ACTB-cre)2Mrt (*hβactin-cre)* heterozygotes (Lewandoski et al. 1997) (*hβactin-cre* breeders were obtained from The Jackson Laboratory, Stock No. 003376). The *Gal^cKO^* and *Dbh^cre^* strains are both available at The Jackson Laboratory as Stock No. 034319 and 033951, respectively. For long-range PCR to identify homologous recombinants for *Gal^cKO^* allele generation, the primer pairs were 5’-GAAGTCCGACAGCTGGTCCATCTGAG and 5’-CACAACGGGTTCTTCTGTTAGTCC for the upstream junction, and 5’-CACACCTCCCCCTGAACCTGAAAC and 5’-GCTAGAAGGATGCTGTATAGAGTAGGCTTC for the downstream junction.

**Figure 1.**
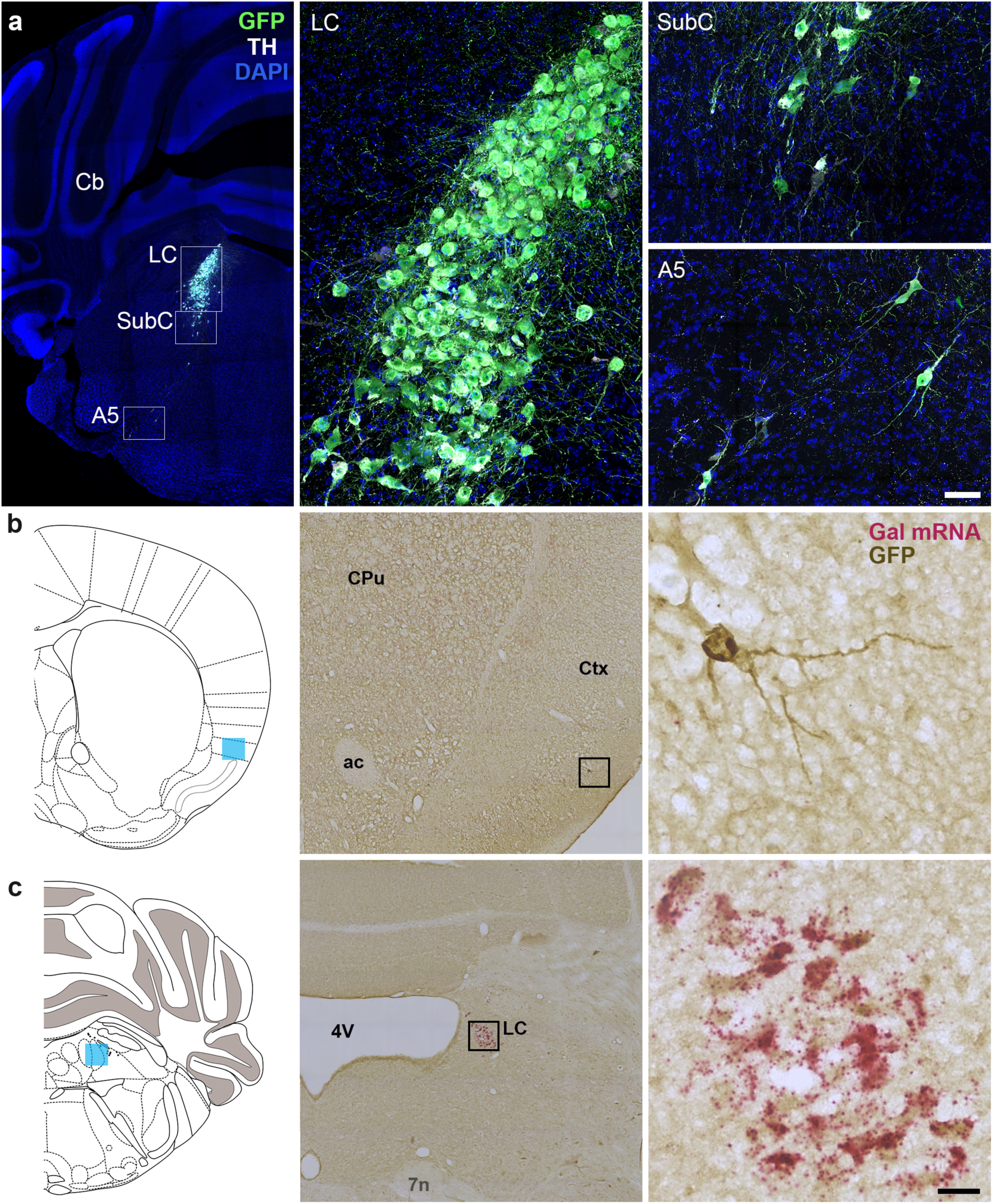
*Dbh^cre^* drives cre recombinase expression in noradrenergic neurons and a sparse population of Gal-negative cortical neurons. (a) Representative coronal sections of *Dbh^cre^;RC::LTG* brains stained for EGFP and TH-expressing neurons of the LC, SubC, and A5. Scale, 379 µm (brain), 50 µm (LC). Cb, cerebellum. LC, locus coeruleus. SubC, sub-coeruleus. (b-c) Coronal sections of *Dbh^cre^; RC::LTG* brains stained for Gal (riboprobe, magenta) and EGFP (DAB antibody, brown). Scattered EGFP+, Gal-negative neurons are observed in the adult cortex (b), while EGFP+ neurons in the LC are Gal-positive (c). Coronal schematics of the adult brain (left) indicate the position of imaged neurons (blue shading). Scale, 20 µm (high magnification), 252 µm (low magnification). ac, anterior commissure. CPu, caudate putamen. Ctx, cortex. LC, locus coeruleus. 4V, fourth ventricle. 7N, facial nerve.

### Stereotaxic delivery of colchicine

Colchicine, a disrupter of microtubule polymerization, was used to amplify galanin peptide signal for experiments requiring immunofluorescent cell body detection, as previously described (Perez et al. 2001; Skofitsch and Jacobowitz 1985). Mice received intracerebroventricular injections of colchicine (10 μg in 0.5 μl sterile saline, 100 nL/min delivery rate) using the following stereotaxic coordinates: 0.48 posterior to bregma, 1.0 mm lateral to midline, and 2.8 mm ventral to skull surface. To treat colchicine-induce sickness, mice were administered acetaminophen (1.6 mg/ml) in the drinking water for 1-2 days before and after surgery. Mice were euthanized for tissue collection approximately 48-hrs following colchicine injection.

### Tissue collection

Adult male and female mice were deeply anesthetized with sodium pentobarbital (0.1 mL of 50 mg/mL i.p.) and perfused transcardially with PBS followed by 4% PFA in PBS. Brains were postfixed overnight by immersion in 4% PFA at 4°C. Following rinse in PBS, tissue was cryoprotected in 30% sucrose in PBS and embedded in Tissue Freezing Medium (General Data Healthcare, Cincinnati, OH). For immunohistochemistry, 40-µm free-floating tissue cryosections were collected on a Leica CM3050-S cryostat (Leica Biosystems, Buffalo Grove, IL). Tissue for *in situ* hybridization was cryosectioned at 14-μm and collected onto Superfrost Plus slides, air dried and stored at −80°C.

To assess Fos activation in the LC, mice were subjected to 1 h of acute restraint stress. Ninety min after the start of the stress, mice were anesthetized with isoflurane and transcardially perfused with potassium phosphate-buffered saline (KPBS) followed by 4% PFA in PBS. Brains were postfixed overnight by immersion in 4% PFA at 4°C, and then transferred to 30% sucrose in KPBS for 48 h at 4°C. Brains were flash frozen in isopentane on dry ice and embedded in Tissue Freezing medium. Tissue for immunohistochemistry was cryosectioned at 40-µm.

Half of the mice used for HPLC were subjected to foot shock stress for 20 min (nineteen 1 mA shocks lasting 0.5 ms with a random intershock interval of 30, 60 or 90 s), and brains were collected 15 min after the end of the foot shock exposure. For galanin ELISA and HPLC, mice were deeply anesthetized with isoflurane and rapidly decapitated for brain extraction. Hypothalamus, PFC, dorsal and ventral hippocampus, pons, and midbrain (ELISA) or PFC, pons, and whole hippocampus (HPLC) were rapidly dissected and snap-frozen in isopentane on dry ice. The samples were weighed and stored at −80°C until processing.

### Immunohistochemistry

Immunohistochemistry was performed as previously described (Sciolino et al. 2016; Robertson et al. 2013). For immunofluorescent staining, noradrenergic cell bodies were labeled with Rabbit anti-TH (1:1000) and Goat anti-rabbit 488 secondary antibody (1:1000) or Goat anti-rabbit 633 (1:1000). Galanin fibers were labeled using rabbit anti-galanin (1:2000) and goat anti-rabbit 488 secondary antibody (1:1000). EGFP-expressing neurons and axons were detected using Chicken anti-GFP primary antibody (1:10,000) and Goat anti-chicken Alexa Fluor 488 secondary antibody (1:1000). For immunoperoxidase staining, we used Chicken anti-GFP primary antibody and biotinylated Goat anti-chicken secondary antibody (1:500), together with Vectastain Elite ABC kit (PK6100), and DAB (SK4100) (all Vector Laboratories). Coverslips were applied using Vectashield hard-set mounting medium with or without DAPI (H-1500 or H-1400, Vector Labs, Burlingame, CA) or Prolong Diamond Anti-Fade mounting medium (P36970, Invitrogen).

Activated neurons were detected with rabbit anti c-fos primary antibody (1:5,000) and goat anti-rabbit 488 secondary (1:600). TH-expressing cells were detected using chicken anti-TH (1:1000) and goat anti-chicken 568 secondary (1:600). After staining, sections were mounted on slides and cover slipped with Fluoromount plus DAPI (Southern Biotech, Birmingham, AL). Antibodies are summarized in **Table 1**.

**Table 1.**
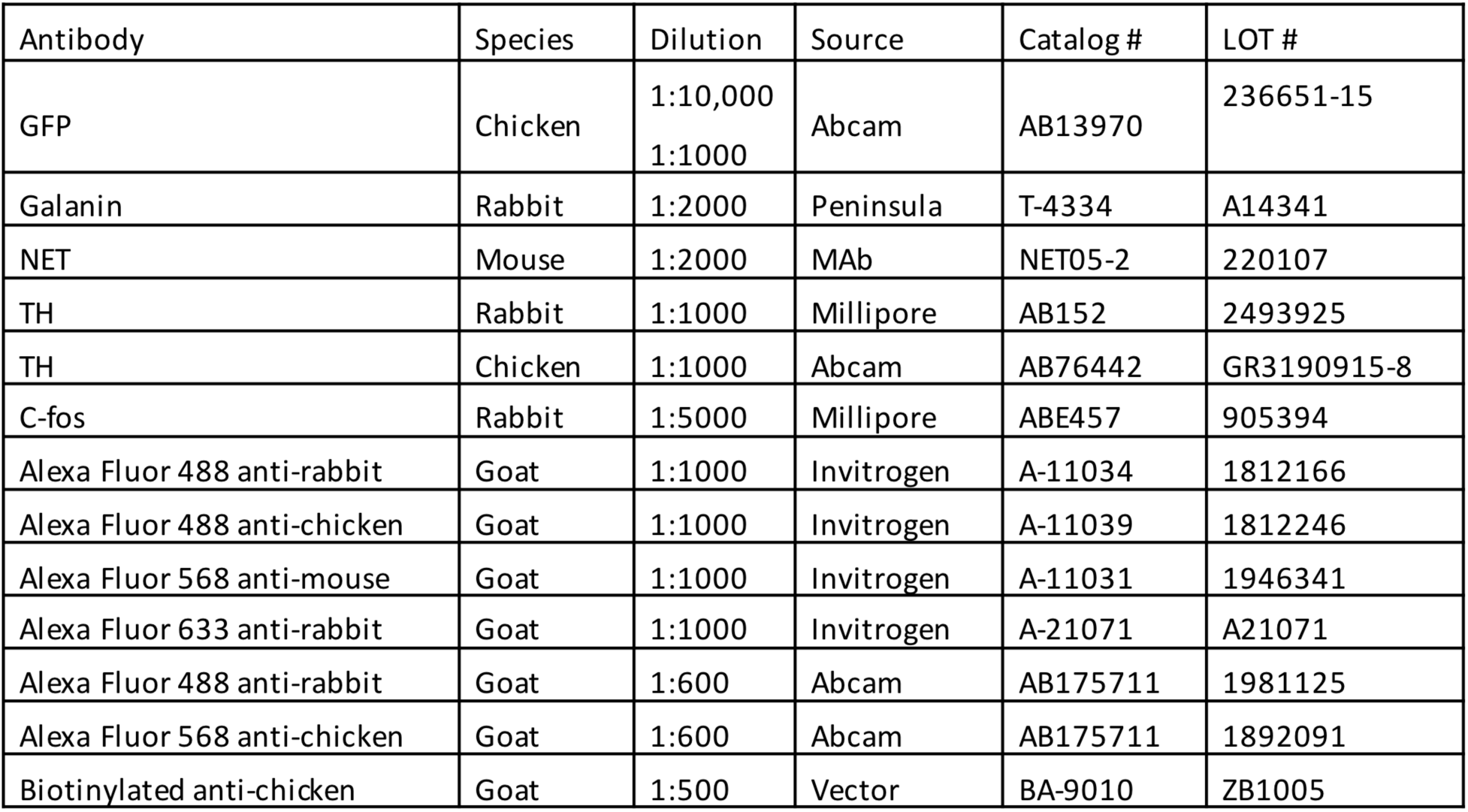
Antibodies.

### *In situ* hybridization

*In situ* hybridization was performed using BaseScope^TM^ Probe BA-Mm-Gal-E3-1ZZ-st (Catalog #712941, Lot #18067A; Advanced Cell Diagnostics, Newark CA), which targets nucleotides 228-266 of mouse *Gal* (exon 3). Tissue was labeled according to the manufacturer’s instructions and then processed for immunohistochemistry as described above.

### Image acquisition

Images of fluorescently-labeled cell bodies were collected on a Zeiss LSM 710 or 880 confocal microscope (Carl Zeiss Microscopy, Thorwood NY) at 10 and 40x. Images of Fos immunofluorescent-labeled sections were collected on a Leica DM6000B epifluorescent upright microscope at 20x. Images of immunoperoxidase-labeled cell bodies were collected on a Leica epifluorescent Zeiss Observer Z1 microscope at 40x. Anatomical location was confirmed by reference to a mouse brain atlas (Paxinos and Franklin 2013).

Images of fluorescently-labeled fibers were collected at 20x on a Zeiss LSM 880 confocal microscope. Three matched sections (425 x 425 μm) were acquired from each subject within the following defined regions relative to bregma: cingulate regions of the PFC A25 and A24a (1.98 to 1.42 mm), hippocampal dentate gyrus (DG) (−1.06 to −1.94), hypothalamic PVN and LHA (−0.58 to −1.22 mm), and midbrain VTA (−2.80 to −3.16) with reference to a mouse brain atlas. In a few cases (<5% of the total samples), two sections instead of three were obtained due to tissue damage or loss.

### Image processing and quantification

For images of immunoperoxidase-labeled GFP-positive cell bodies, 8-μm Z-stacks were converted to a single image using the extended focus feature of Zen 2012 Blue Software (Carl Zeiss). For images of immunofluorescent-labeled galanin-positive cell bodies, a colocalization mask was created in the MIP image. Mask thresholds were empirically determined and applied equally to the control and mutant groups using NIH Image J (https://imagej.nih.gov/ij/). Using an ‘AND’ operation for co-expression between galanin and TH, the masks represented pixels with no expression (black) and co-expression (white). To optimize the full dynamic range of the fluorescence signal, only brightness and contrast adjustments were applied.

For quantification of Fos immunofluorescence, three LC sections from each mouse were analyzed and averaged together to determine group means. All image analysis was performed with ImageJ software for background subtraction, threshold application (Otsu method) and quantification based on size and shape criteria for Fos positive nuclei (30–70 mm^2^, circularity 0.7–1.0). TH immunoreactivity was used to define LC area and only neurons within that area were used for Fos quantification. Experimenter was blind to genotype during image collection and analysis.

For quantification of galanin fibers, we first removed fluorescent artifacts from our 425 x 425 µm^2^ image using the DEFiNE macro (https://figshare.com/s/1be5a1e77c4d4431769a) ‘Clean Images’ function with default settings to subtract artifacts, both large and small, that were morphologically distinct from fibers, as previously described (Powell et al. 2018). To remove blood vessels from the maximum intensity projection (MIP) images, we then performed two subtractions to remove particles larger than 65 µm^2^ by converting the images to binary format at a threshold of 1 standard deviation above average pixel intensity. Large artifacts were identified using FIJI’s ‘Analyze Particle’ function and were removed from the grayscale MIP using FIJI’s ‘Image Calculator – AND’ function.

An experimenter blind to treatment group used DEFiNE’s ‘Quantify Fiber’ function to set background-sensitive thresholds for the single-channel z-stack image. FIJI’s ‘Skeletonize’ function was then applied to the resulting binary images to normalize variability in axon circumference. The number of µm^2^ pixels in each binary image was measured for the galanin channel. For each subject, the number of pixels in the images from each region was averaged. Representative images were matched for anatomy and were converted to grayscale to enhance visibility of the fibers.

### Galanin Enzyme-linked Immunosorbent Assay (ELISA)

Samples were put in buffer (2.5% aprotinin in 0.5M acetic acid), homogenized, heated (100°C for 10 min), and centrifuged (30 min at 4°C, 3000 rpm). Supernatant was collected, evaporated in a vacuum-sealed concentrator (10 h at 60°C and 20,000mm Hg; Labconco Centrivap) and then stored at −20°C. Samples were reconstituted in 250 μL EIA buffer (Peninsula Laboratories, San Carlos, CA), and processed according to the manufacturer’s instructions (Galanin Rat and Mouse ELISA kit, S1208, Peninsula Laboratories, San Carlos, CA). Wells were read at 450 nm, averaged across duplicates, and a curve of best fit was used to calibrate to standards.

### High Performance Liquid Chromatography (HPLC)

Tissue was thawed on ice and sonicated at 4°C in 0.1 N perchloric acid (10 μl/mg tissue) for 12 sec of 0.5 sec pulses. Sonicated samples were centrifuged (16100 rcf) for 30 min at 4°C, and the supernatant was then centrifuged through 0.45 µm filters at 4000 rcf for 10 min at 4°C. For HPLC, an ESA 5600A CoulArray detection system, equipped with an ESA Model 584 pump and an ESA 542 refrigerated autosampler was used. Separations were performed at 23°C using an MD-150 × 3.2 mm C18 column. The mobile phase consisted of 90 mM sodium acetate, 37 mM citric acid, 0.2 mM EDTA, 0.4 mM 1-octanesulfonic acid sodium, 0.025% triethylamine, and 4.5% methanol at pH 4.4. A 25 µl of sample was injected. The samples were eluted isocratically at 0.4 mL/min and detected using a 6210 electrochemical cell (ESA, Bedford, MA) equipped with 5020 guard cell. Guard cell potential was set at 600 mV, while analytical cell potentials were −175, 150 and 400 mV. NE and its primary metabolite MHPG were measured with electrochemical detection. Analytes were identified by matching criteria of retention time and sensor ratio measures to known standards (Sigma-Aldrich, St. Louis, MO). Compounds were quantified by comparing peak areas to those of standards on the dominant sensor.

### Behavior

Adult male and female mice were tested between 3 and 8 months of age, and sexes were balanced for all tests. Mice were group-housed for the duration of behavioral testing, unless stated otherwise. Tests were separated by at least 4 days.

#### Circadian locomotor activity

Mice were placed in automated locomotor recording chambers (transparent Plexiglas cages on a rack; San Diego Instruments, La Jolla, CA), and ambulation (consecutive photobeam breaks) were recorded for 23 h in 30 min time bins.

#### Fear conditioning

Fear conditioning training, contextual fear testing, and cued fear testing were conducted over 3 consecutive days. The chamber (Coulbourn Instruments, Holliston, MA) was equipped with a house light, ceiling-mounted camera, speaker and an electric grid shock floor that could be replaced with a non-shock wire mesh floor. Chambers were cleaned in between animals with Virkon. The acquisition trial on day 1 lasted 6 min, followed by 3 tone-shock pairings during which the tone was present for 20 s and was co-terminated with a 3 s, 0.5 mA foot shock. Mouse behavior was recorded for 60 s following tone-shock presentation before the next round. Contextual fear testing on day 2 was performed in the same chamber as day 1 and lasted 7 min without any presentation of tone or shock. Cued fear testing on day 3 was conducted in a different chamber than days 1 and 2, with a non-shock mesh floor instead of the previous shock grid floor. The cued fear testing trial lasted 8 min, with the tone starting after 3 min and continuing until the end of the trial. Trials were programmed and run using the FreezeFrame software (Coulbourn) to automatically record freezing behavior during each trial.

#### Forced swim test

Mice were placed in a 3.5 L beaker with 3 L of water (25°C) and behavior was videotaped for 6 min. Behavior in the last 4 min of the test was scored by an observer blind to genotype with immobility defined as lack of any movements besides those which are required for balance and to keep the animal’s head above water.

#### Tail suspension test

Climbstoppers (1.5 in long, clear flexible tubing) were placed over the base of each animal’s tail to prevent tail climbing (Can et al. 2012). A piece of medical tape was placed around the tail, halfway between the base and the tip, and the mouse was suspended from a ring stand 16-18” from the counter. Behavior was videotaped for 6 min and the entire test was later scored by an observer blind to genotype, with immobility defined as a lack of all movement.

#### Sucrose preference test

Mice were individually housed at least 1 week prior to the start of sucrose preference testing. Two small water bottles (modified 15mL conical tubes), one filled with 1% sucrose water and the other with tap water, were placed in the home cage. Bottles were weighed every 24 h, refilled as necessary, and the location of the bottles was switched every 24 h to control for side bias. The amount of water consumed from the bottles during each 24-h period was calculated by subtracting the weight from the previous measurement. Percent sucrose preference was then calculated as (sucrose water consumed) ÷ (normal + sucrose water consumed) × 100.

#### Open field

Mice were placed in the open field arena (36” x 36” x 36”) for 5 min under dim light. An overhead camera and TopScan software were used to automatically detect the time spent in the center of the square, total distance travel, and ambulation velocity.

#### Zero maze

Mice were placed in the zero maze (2” wide track, 20” diameter) for 5 min under dim light. An overhead camera and TopScan software (Clever Sys Inc., Reston, VA) were used to automatically detect the time spent in open arms.

#### Elevated plus maze

Mice were placed in the elevated plus maze (25” by 25” arms, 2.5” track width) for 5 min under dim light. An overhead camera and TopScan software were used automatically detect the time spent in open arms.

#### Novelty-suppressed feeding

Following 24-h food deprivation, mice were placed in the corner of a novel arena (36” x 36” x 36”) that was illuminated by red light and contained a regular chow pellet in the center of the arena. Each food pellet was weighed before the start of the experiment. Mice were observed for up to 10 min while the latency to initiate feeding behavior (biting the food) was recorded. If the mouse did not bite the pellet after 10 min elapsed, the experiment ended, and a value of 600 s was assigned. To control for appetite, each animal was moved individually to a new home cage that contained the food pellet from the previous arena, and food intake was measured for 1 h. The arena was cleaned with Virkon between animals.

#### Marble burying

Twenty black glass marbles were arranged in a grid pattern on top of bedding (2” deep) in a cage (10” x 18” x 10”) with no lid. The mouse was placed in the center of the cage and left undisturbed for 30 min under bright light. At the end of the test period, images of each cage were taken from the same distance and angle. The number of marbles buried was determined by an experimenter blind to genotype by counting the number of marbles that remained unburied and subtracting that count from 20. Marbles were considered buried when at least 2/3 of the marble could not be seen.

#### Shock probe defensive burying

The test cage (13” x 7” x 6”) was filled with approximately 3 cm of clean bedding, and a shock probe was located on one wall of a chamber (14 cm long and 0.5 cm in diameter). The probe extended 10 cm into the cage and was wrapped with 2 alternating copper wires that were connected to a shock generator (Coulbourne). For testing, mice were placed in the cage at the opposite side from the probe. When the animal touched the probe, they received a 0.5 mA shock, and behavior was videotaped for 15 min from the time of the first shock (Degroot and Nomikos 2004). The probe remained electrified throughout the test. Videos were later scored by an observer blind to genotype. Measured behaviors included number of probe touches, time spent digging, freezing, grooming, and number of rearing events.

#### Nestlet shredding

Mice were individually placed in novel cages (13” x 7” x 6”) with 1 cm of clean bedding and a cotton fiber nestlet (5 cm x 5 cm, 0.5 cm thick, ∼2.5 g) placed on top of the bedding. Each nestlet was weighed prior to the start of testing. Mice were left undisturbed for 2 h, after which the weight of the remaining untorn nestlet was recorded and the percent shredded by each mouse was calculated.

### Statistical Analysis

Data were found to be normally distributed using the D’Agostino-Pearson test. Data were analyzed via unpaired t-test, one-way or two-way ANOVA with *post hoc* Tukey’s test for multiple comparisons when appropriate. Significance was set at p<0.05 and two-tailed variants of tests were used throughout. Data are presented as mean ± SEM. Calculations were performed and figures created using Prism Version 7 and 8 (GraphPad Software, San Diego, CA).

## RESULTS

### *Gal^cKO-Dbh^* mice lack galanin mRNA and protein in noradrenergic neurons

To characterize the role of galanin in noradrenergic neurons, we first generated a knock-in allele that expresses cre recombinase under control of the noradrenergic-specific dopamine **β**-hydroxylase promoter (*Dbh^cre^*). To evaluate the expression pattern and activity of *Dbh^cre^*, we crossed it to the cre-responsive reporter *RC::LTG* (Plummer et al. 2017) which drives EGFP expression upon *cre* recombination. We observed EGFP expression in tyrosine hydroxylase+ (TH) noradrenergic neurons in the peripheral and central nervous system of *Dbh^cre^; RC::LTG* double heterozygotes (**Fig. 1a** and data not shown). Similar to some *Dbh-cre* transgenic alleles (Matsushita et al. 2004; Gerfen et al. 2013), we also observed scattered EGFP+ cells in the adult cortex that likely reflect early transient expression of *Dbh* (**Fig. 1**). Importantly for the present study, the EGFP+ cortical neurons were negative for *Gal* mRNA expression (**Fig. 1**).

We next generated a conditional knockout allele of *Gal* (*Gal^cKO^*) and crossed it with *Dbh^cre^* to create the experimental mice referred to as *Gal^cKO-Dbh^* mutants (**Fig. 2**). To confirm that mutants lack galanin expression selectively in noradrenergic neurons, we used *in situ* hybridization to detect *Gal* mRNA in adult brains of *Gal^cKO-Dbh^* mutant and littermate control mice*. Gal* mRNA was abundant in LC and SubC noradrenergic neurons of control mice, but completely absent in these nuclei in *Gal^cKO-Dbh^* mutants (**Fig. 2**). Importantly, *Gal* mRNA expression remained intact in TH-negative neurons in C2/A2 and other non-noradrenergic brain regions, including the bed nucleus of the stria terminalis (BNST), lateral preoptic area, lateral hypothalamus and inferior olive (**Fig. 2 and 3**).

**Figure 2.**
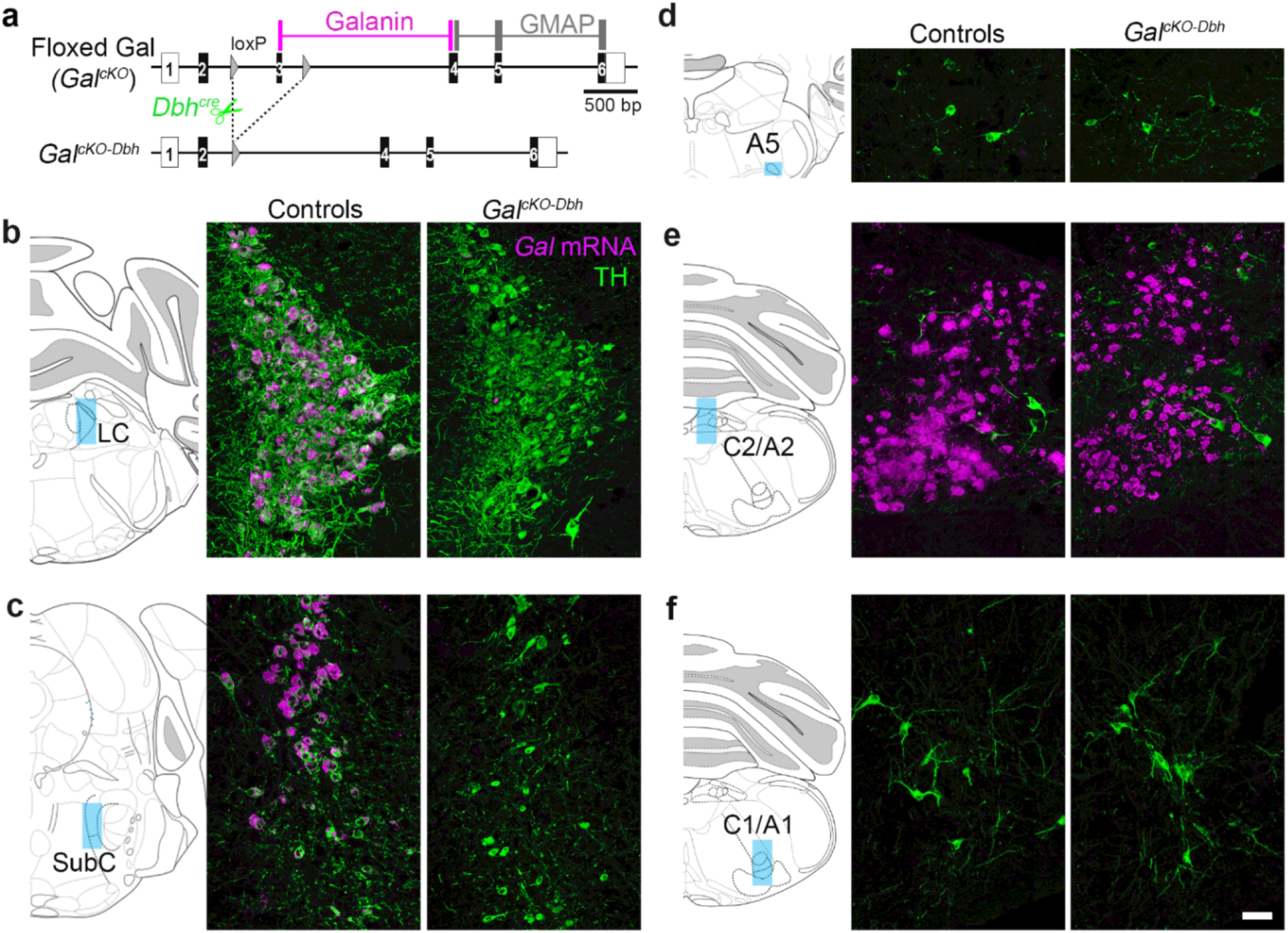
*Gal* conditional knockout allele permits selective disruption of galanin synthesis in noradrenergic neurons. (a) Schematic diagram of *Gal^cKO^* allele. Recombination by *Dbh^cre^* leads to disruption of galanin synthesis in NE neurons. (b-f) *Left*. Schematic illustration of coronal mouse brain showing location of images. *Right*. Representative coronal brain sections stained for *Gal* mRNA (riboprobe; magenta) and tyrosine hydroxylase (TH antibody; green). *Gal* mRNA is absent from the SubC and LC of mutant mice (*Gal^cKO-Dbh^*). Scale bar, 50 µm. LC, locus coeruleus. SubC, sub-coeruleus.

**Figure 3.**
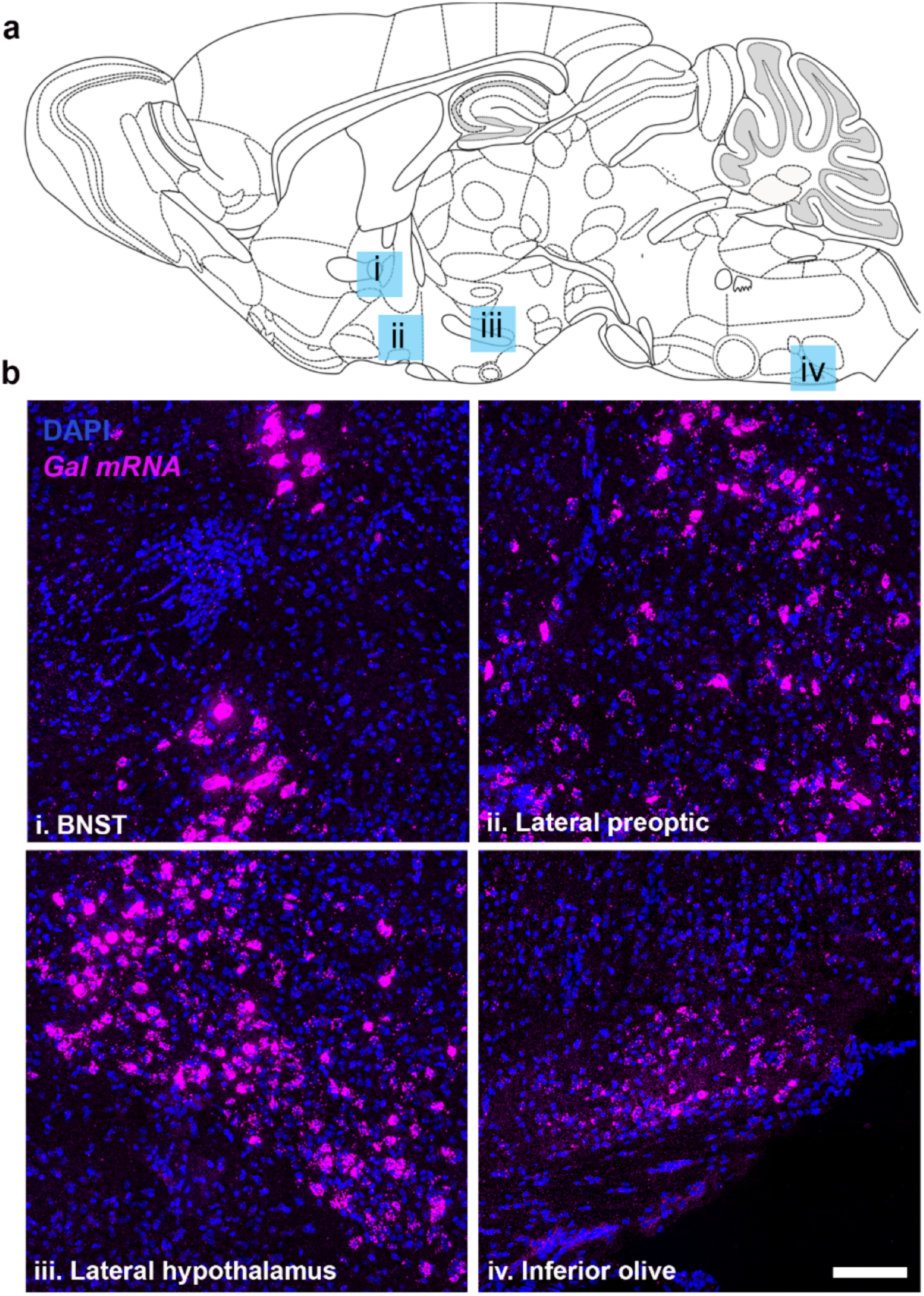
*Gal* mRNA expression persists outside the noradrenergic system of *Gal^cKO-Dbh^* mice. (a) Schematic illustration of sagittal mouse brain showing location of images. (b) Representative sagittal sections from brains of *Gal^cKO-Dbh^* mice stained for *Gal* mRNA (riboprobe; magenta) and DAPI (blue). Scale, 100 µm. BNST, bed nucleus of the stria terminalis.

Next, we examined galanin protein expression by immunohistochemistry. To induce accumulation of the peptide in neuronal somata and thus improve its visualization, we injected the axonal transport blocker colchicine (10 μg) into the lateral ventricle of adult *Gal^cKO-Dbh^* mutant and littermate control mice. Like the pattern of mRNA expression, galanin was evident in TH+ neurons of the LC and SubC of littermate controls but abolished in almost all noradrenergic neurons in *Gal^cKO-Dbh^* mutants (**Fig. 4** and data not shown). In both *Gal^cKO-Dbh^* mutant and littermate control mice, we observed galanin immunoreactivity in non-noradrenergic populations, including the inferior olive (**Fig. 4** and data not shown). Interestingly, galaninergic fibers persisted in the LC region of *Gal^cKO-Dbh^* animals (**Fig. 4**), revealing that LC cells are innervated by galanin-containing neurons from non-noradrenergic sources. Combined, these findings indicate that *Gal^cKO-Dbh^* mice have a selective disruption of galanin synthesis in noradrenergic neurons.

**Figure 4.**
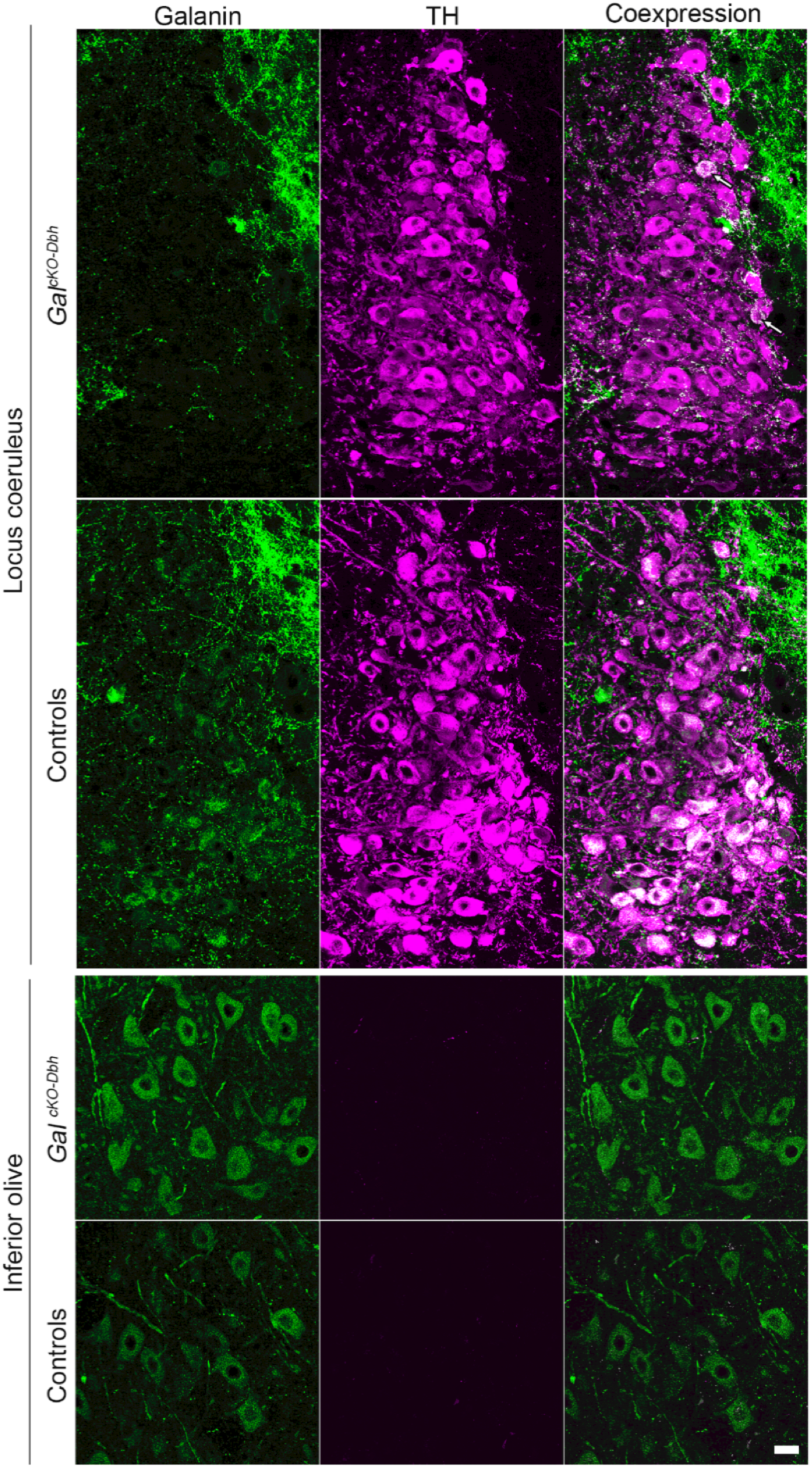
Galanin protein expression is reduced selectively in noradrenergic neurons of *Gal^cKO-Dbh^* mice. Representative coronal brain sections stained for galanin (antibody; green) and tyrosine hydroxylase (TH antibody; purple). *Top*. In littermate controls, the majority of noradrenergic locus coeruleus (LC) neurons co-express TH and galanin (white). In *Gal^cKO-Dbh^* mice, galanin (green) is absent from the majority of TH+ (purple) LC neurons. Arrows indicate an occasional TH+ neuron in in *Gal^cKO-Dbh^* mice that co-expresses galanin (white). Dense TH-negative, galanin+ fibers (green) surround and innervate the LC in *Gal^cKO-Dbh^* mice and littermate controls. *Bottom*. Galanin is expressed normally in non-noradrenergic olivary neurons of *Gal^cKO-Dbh^* mice and littermate controls. Scale, 20 µm.

### Noradrenergic neurons provide a significant proportion of galanin to select brain regions

To assess the proportion of galanin in the brain that comes from noradrenergic neurons, we measured galanin protein levels in dissected tissue from the prefrontal cortex (PFC), dorsal and ventral hippocampus, hypothalamus, midbrain, and pons by ELISA. As a negative control and for comparison, we also generated and tested mice lacking galanin throughout the brain (*Gal^cKO-nestin^*). *Gal^cKO-Dbh^* mice had significantly decreased galanin in the PFC (*t*_8_=3.069, p=0.0154), pons (*t*_9_=2.823, p=0.020), dorsal hippocampus (*t*_8_=6.593, p=0.0002), and ventral hippocampus (*t*_8_=2.314, p=0.0494) compared to littermate controls, but no difference in the midbrain (*t*_7_=0.5440, p=0.6033) or hypothalamus (*t*_10_=0.4800, p=0.6415) (**Fig. 5**). These findings suggest that noradrenergic neurons provide a significant source of galanin to the cortex and hippocampus.

**Figure 5.**
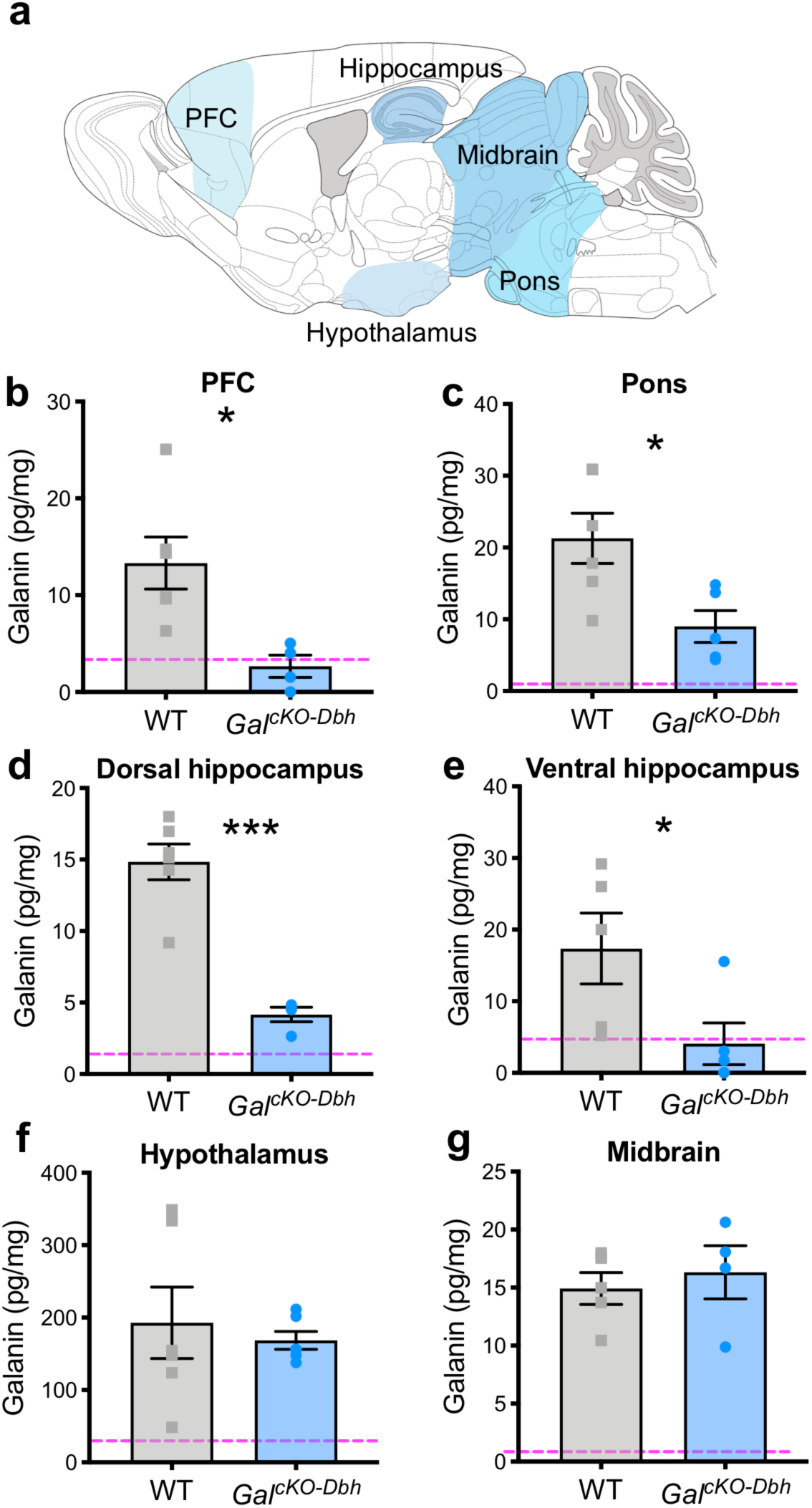
*Gal^cKO-Dbh^* mice have decreased galanin in the hippocampus, prefrontal cortex, and pons. Tissue galanin levels were measured by galanin ELISA. Tissue was collected via rapid dissection of whole regions (see methods for more detail) (a). *Gal^cKO-Dbh^* mice showed significantly decreased galanin in the PFC (b), pons (c), and dorsal and ventral hippocampus (d, e), with no differences in the hypothalamus (f) or midbrain (g) compared to littermate controls. Dashed line shows the average signal determined in each discrete brain region of negative control *Gal*^cKO-nestin^ tissue. In the PFC and hippocampus, *Gal^cKO-Dbh^* galanin levels were reduced to an equivalent level as seen in the negative control samples (a, c, d). *n* = 4-6 mice per group. Data were analyzed by independent samples t-tests (two-tailed). Error bars show SEM. Error bars show SEM. *p<0.05, ***p<0.001

To provide finer neuroanatomical resolution, we quantified the density of galanin-positive fibers in the cingulate cortex (A25 and A24a), hippocampal dentate gyrus (DG), lateral and paraventricular nuclei of the hypothalamus (LHA and PVN) and midbrain ventral tegmental area (VTA) of *Gal^cKO-Dbh^*, littermate control, and galanin null mice (*Gal^NULL^*). We observed a significant reduction in galanin fiber expression in the A25 (*t*_13_=2.276, p=0.0404), A24a (*t*_13_=2.283, p=0.0399) and DG (*t*_17_=2.536, p=0.0213) brain areas of *Gal^cKO-Dbh^* mice (**Fig. 6**), revealing these discrete regions within the prefrontal cortex and hippocampus are innervated by noradrenergic-derived galanin neurons. By contrast, *Gal^cKO-Dbh^* and littermate controls exhibited similar levels of galanin fiber expression in the midbrain VTA (*t*_16_=0.7647, p=0.4556) as well as the hypothalamic PVN (*t*_17_=0.7152, p=0.4842) and LHA (*t*_17_=0.7952, p=0.4375) (**Fig. 6**), demonstrating that these areas do not receive significant innervation by noradrenergic-derived galanin neurons.

**Figure 6.**
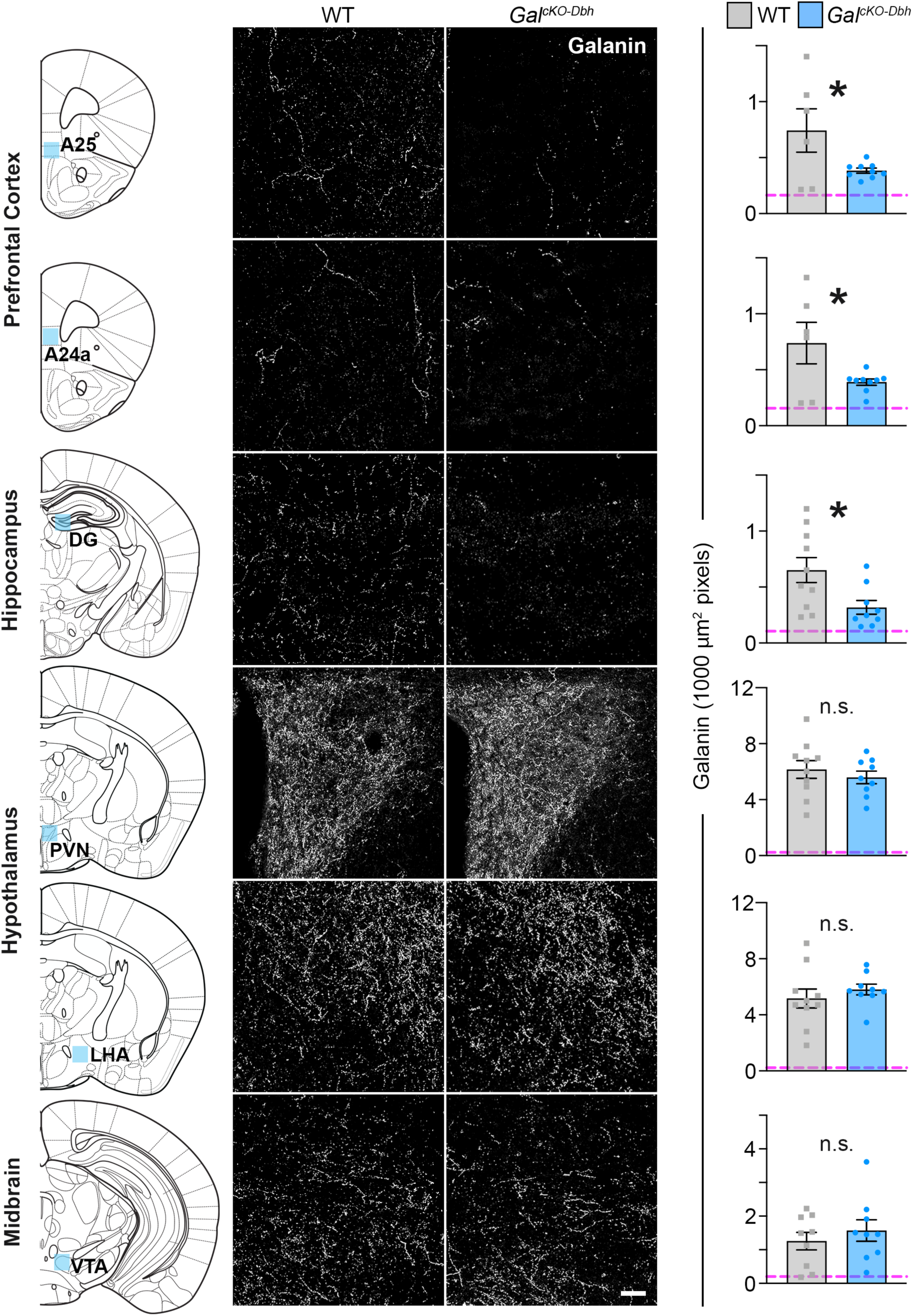
*Gal^cKO-Dbh^* mice have a reduction in galanin-positive fibers in subregions of the prefrontal cortex and hippocampal dentate gyrus. *Left*. Blue shading on schematic of coronal mouse brain shows the location of fiber quantification. *Middle*. Representative coronal brain sections immunostained for galanin (white). Scale bar, 50 µm. *Right*. Average galanin fiber density in discrete brain regions. Dashed line shows the average background signal determined in each discrete brain region of *Gal^NULL^* tissue. *Gal^cKO-Dbh^* mice showed reduced galanin fiber expression in prefrontal cortex (A25 and A24a) and hippocampus (DG), with no differences in the hypothalamus (PVN and LHA) or midbrain (VTA) compared to littermate controls. *n* = 6-10 littermate controls, *n* = 9 *Gal^cKO-Dbh^* mutants. Data were analyzed by independent samples t-tests (two-tailed). Error bars show SEM. *p<0.05. n.s., non-significant.

### Normal NE abundance and turnover in *Gal^cKO-Dbh^* mutants

To determine whether loss of noradrenergic-derived galanin affects NE metabolic activity, we measured concentrations of NE, its major metabolite MHPG (3-Methoxy-4-hydroxyphenylglycol), and turnover ratio of MHPG:NE in dissected tissue from the PFC, pons, and whole hippocampus by HPLC at baseline and immediately after foot shock stress (**Fig. 7**). A two-way ANOVA showed that stress caused a significant decrease in NE in the pons (*F*_1,20_=7.282, p=0.014) and hippocampus (*F*_1,20_=5.653, p=0.028), but not PFC (*F*_1,20_=0.255, p=0.619), and a significant increase in MHPG in all three regions (pons, *F*_1,20_=37.27, p<0.0001; hippocampus, *F*_1,20_=27.4, p<0.0001; PFC, *F*_1,20_=11.26, p<0.01). This resulted in an overall increase in NE turnover, as measured by the MHPG:NE ratio in the pons (*F*_1,20_=698.9, p<0.0001), hippocampus (*F*_1,20_=195.8, p<0.0001), and PFC (*F*_1,20_=23.4, p<0.001). No genotype differences were observed (**Fig. 7a-c**).

**Figure 7.**
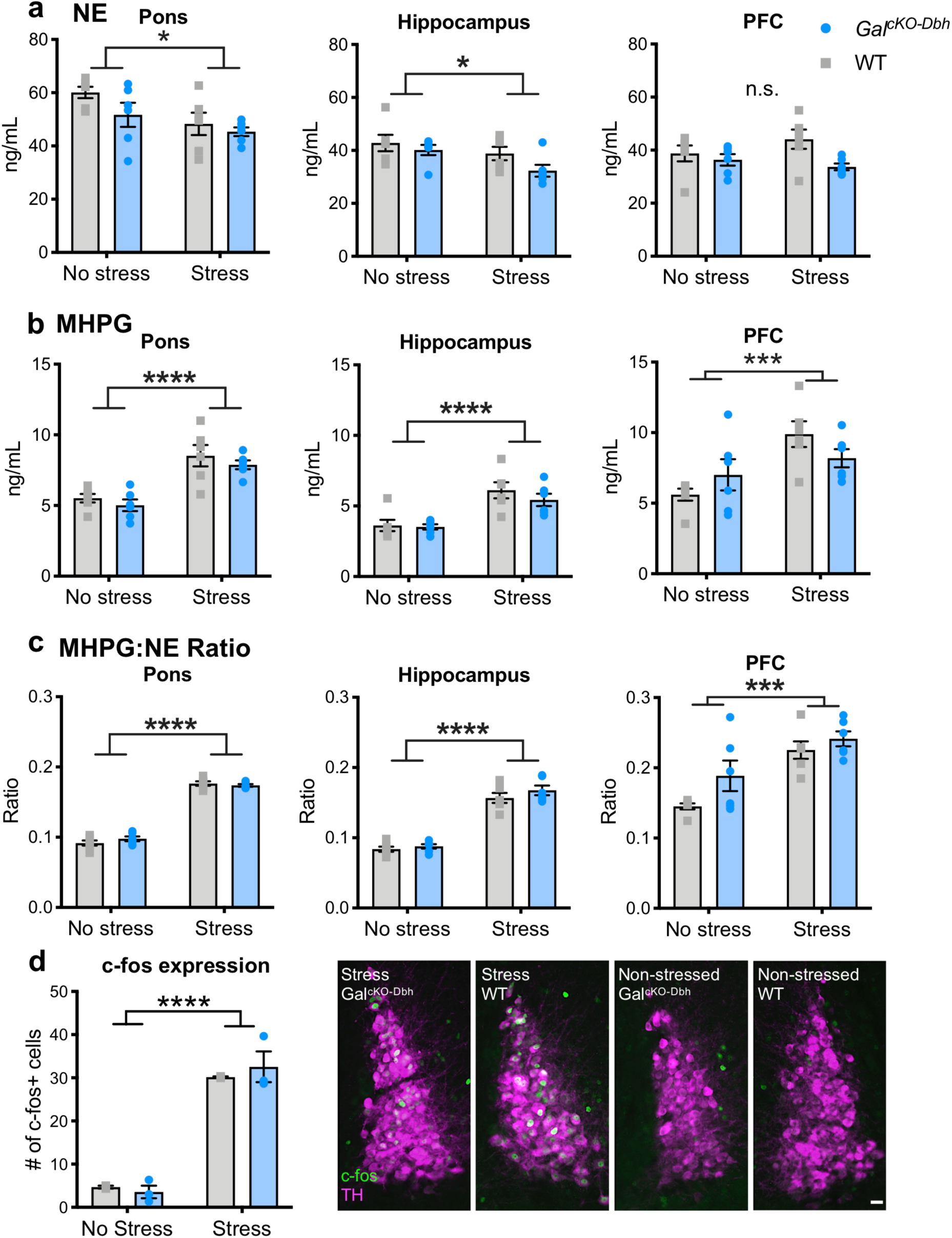
Baseline and stress-induced NE turnover and LC c-fos expression are normal in *Gal^cKO-Dbh^* mice. NE (a) and the major metabolite of NE, MHPG (b), were measured via HPLC at baseline and after acute foot shock stress in the pons, hippocampus, and PFC. Foot shock stress caused an overall decrease in NE and increase in MHPG, leading to an increase in the MHPG:NE ratio in all three brain regions. No differences were seen between genotypes in either the raw measurements or the MHPG:NE ratio. *n* = 6 mice per group. Fos immunohistochemistry and quantification did not show any differences between *Gal*^cKO-Dbh^ mice and WT littermate control mice at baseline or after acute restraint stress (d) *Right*. Representative LC sections stained for Fos (antibody; green) and tyrosine hydroxylase (TH antibody; magenta). *n* = 3 slices per animal, 2-3 animals per group. Data were analyzed by two-way ANOVA. Scale bar, 20 µm. Error bars show SEM. *p<0.05, ***p<0.001, ****p<0.0001, n.s., non-significant.

Given that somatodendritic release of galanin is thought to inhibit LC activity (Vila-Porcile et al. 2009; Xu et al. 2001; Pieribone et al. 1995), we next wanted to determine if LC activity is influenced by the absence of noradrenergic-derived galanin. To do this, we measured Fos, an immediate early gene used as a proxy for neuronal activation, at baseline and after a mild stressor, acute restraint. A two-way ANOVA comparing *Gal^cKO-Dbh^* mutants and controls at baseline and after stress showed that, as expected, stress increased Fos expression in the LC (*F*_1,6_=119.7, p<0.0001). However, there was no genotype effect (*F*_1,6_=0.065, p=0.806), indicating that Fos expression was comparable between *Gal^cKO-Dbh^* mutants and littermate controls (**Fig. 7d**). Together, these findings suggest that NE transmission remains intact when galanin is absent from noradrenergic neurons. It is important to keep in mind, however, that these measures are indirect and lack the sensitivity to detect subtle alterations in noradrenergic system function.

### *Gal^cKO-Dbh^* mice display a more proactive coping strategy in anxiogenic tasks measuring active defensive behaviors

Next, we assessed the behavior of adult *Gal^cKO-Dbh^* mutants and littermate control mice in a battery of anxiety-, depression-, learning-, and motor-related tests. We observed no differences between *Gal^cKO-Dbh^* and littermate control mice in canonical tests of anxiety-like behavior (elevated plus maze, *t*_22_=0.9379, p=0.3585; zero maze, *t*_24_=0.0951, p=0.925; open field, *t*_36_=0.1769, p=0.8606) (**Fig. 8a-c**) or depressive-like behavior (tail suspension test, *t*_23_=1.366, p=0.1853; forced swim test, *t*_23_=1.55, p=0.1347; sucrose preference, *F*_1,16_=0.004, p=0.9479) (**Fig. 8d-f**). In addition, *Gal^cKO-Dbh^* exhibited normal cognitive responses in contextual (*F*_1,20_=0.1053, p=0.7489) and cued fear conditioning (*F*_1,20_=2.778, p=0.1111) (**Fig. 8g**), as well as circadian locomotor activity (*F*_1,22_=0.5295, p=0.4745) (**Fig. 8h**).

**Figure 8.**
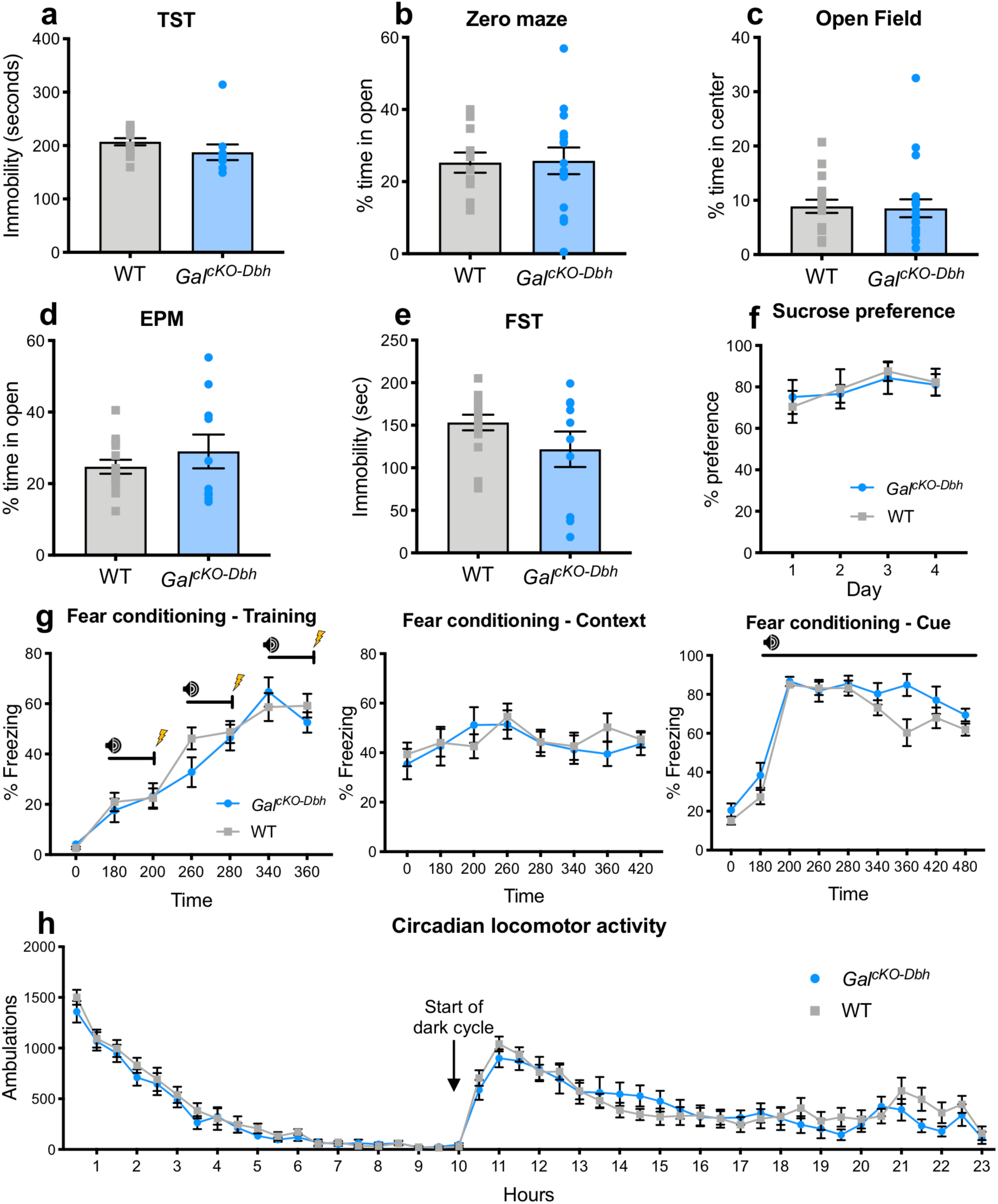
*Gal*^cKO-Dbh^ display normal behavior at baseline in canonical tests for anxiety- and depression-like behavior, locomotion, and associative learning. There were no differences between *Gal*^cKO-Dbh^ and WT littermate control mice in elevated plus maze (EPM) (a), zero maze (b), open field (c), tail suspension test (TST) (d), forced swim test (FST) (e), sucrose preference (f), fear conditioning (g), or locomotor activity (h). *n* = 10-20 per group; mixed males and females. Error bars show SEM.

Having established that *Gal^cKO-Dbh^* mutants preform normally in standard tests of anxiety and depression that are reliant on *passive* behavioral responses (e.g. immobility, avoidance of open spaces, freezing), we next examined non-canonical anxiety tasks that assess *active* coping behaviors (e.g. digging). Using the marble burying assay, we observed that *Gal^cKO-Dbh^* mice buried significantly more marbles than littermate control mice (*t*_36_=3.85, p<0.001) (**Fig. 9a**), indicating an enhancement of proactive defensive behavior. Mutants showed no difference in repetitive/compulsive behavior during the nestlet shredding task (*t*_18_=0.4845, p=0.634) (**Fig. 9b**). Similarly to the marble burying results, in the shock probe defensive burying assay, *Gal^cKO-Dbh^* mice showed increased time spent in active digging (*t*_15_=2.191, p=0.0447) (**Fig. 9c**), and a trend towards a decrease in passive freezing (*t*_15_=1.601, p=0.1303) (**Fig. 9d**). Additional analysis of concurrent exploratory behaviors revealed *Gal^cKO-Dbh^* mice and littermate controls exhibited similar time spent grooming (WT 47.57 ± 13.95, *Gal^cKO-Dbh^* 61.44 ± 16.75; *t*_15_=0.6271, p=0.54) and number of rearing events (WT 35.75 ± 4.515, *Gal^cKO-Dbh^* 36.78 ± 5.444; *t*_15_=0.1431, p=0.8881). There was no difference in the number of times the mice were shocked by the probe (WT 2.375 ± 0.5957, *Gal^cKO-Dbh^* 2.444 ± 0.3379; *t*_15_=0.1044, p=0.9182), indicating that alterations in behavior were not caused by differences in higher numbers of shocks to one group. We next examined performance in the novelty-suppressed feeding task, a conflict test that pits a mouse’s innate fear of an unfamiliar environment against a desire to feed. *Gal^cKO-Dbh^* mice were quicker to start eating (*t*_34_=2.348, p=0.0248) (**Fig. 9e**), but ate a similar overall amount (*t*_20_=1.331, p=0.1983) (**Fig. 9f**), suggesting the *Gal^cKO-Dbh^* phenotype was not driven by increased appetite. Taken together, these data indicate that selective galanin gene inactivation from noradrenergic neurons shifts defensive behavior to an active coping strategy under anxiogenic conditions.

**Figure 9.**
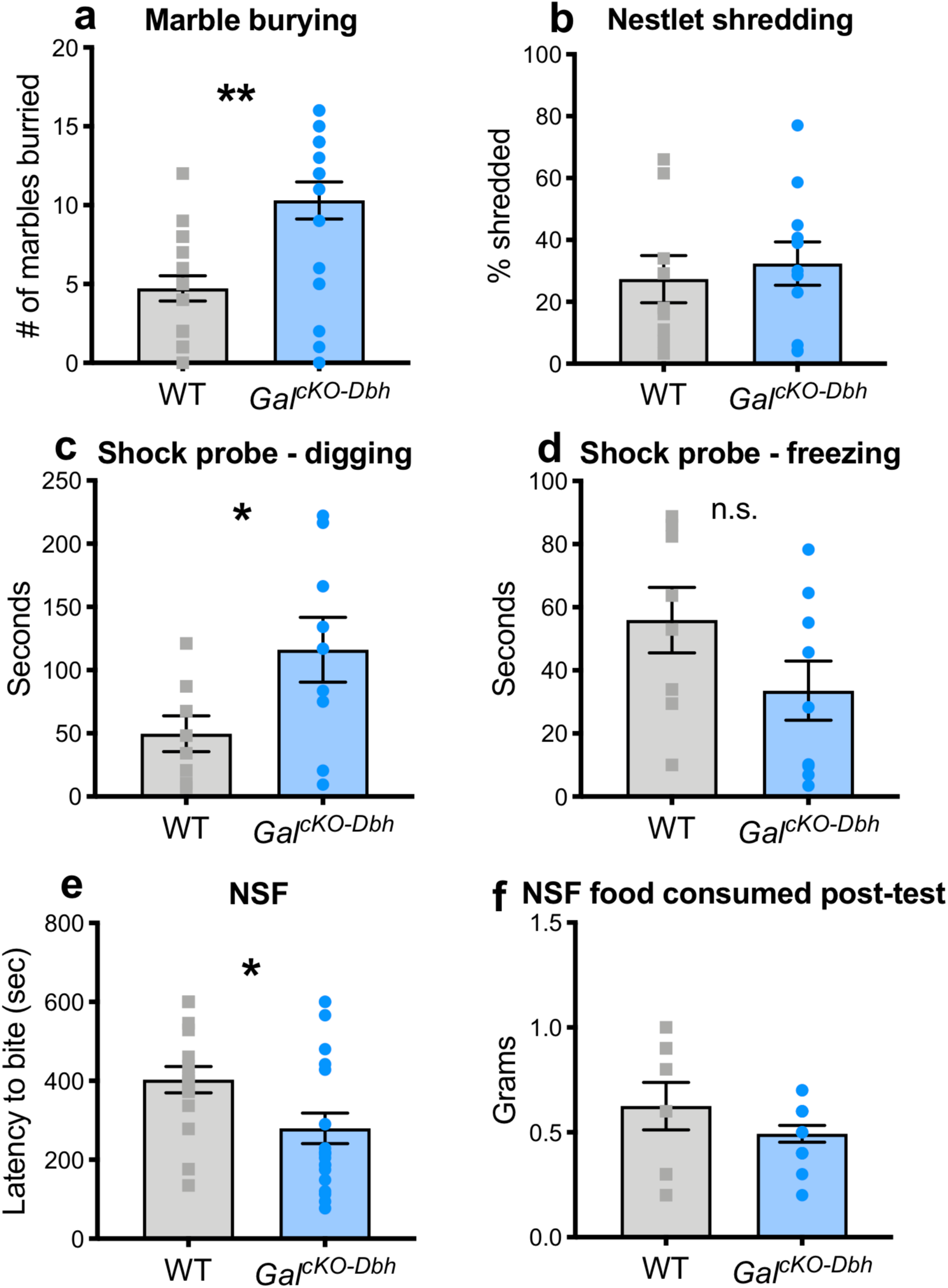
*Gal*^cKO-Dbh^ display increased digging behavior in the marble burying and shock probe defensive burying tasks and decreased latency to feed in the novelty-suppressed feeding task. In the marble burying assay (a), *Gal*^cKO-Dbh^ mice buried significantly more marbles than WT littermate control mice. There were no differences between *Gal*^cKO-Dbh^ and WT controls in nestlet shredding behavior (b). *Gal*^cKO-Dbh^ mice showed increased digging in the shock probe defensive burying task and a trend towards decreased freezing (c, d). In the novelty-suppressed feeding task (NSF), *Gal*^cKO-Dbh^ mice showed a decrease in latency to feed but consumed the same amount of food as WT controls in the hour after the test (e, f). *n* = 10-20 per group; mixed males and females. Error bars show SEM. *p<0.05, ***p<0.001

## DISCUSSION

We have generated the first cell type-specific galanin knockout mouse by selectively deleting the *Gal* gene in noradrenergic neurons, leading to a loss of Gal mRNA and protein in these cells. Our results indicate that a large proportion of the galanin in the hippocampal DG and PFC, as well as about half the galanin in the brainstem, is derived from noradrenergic neurons, whereas alternate sources are responsible for the majority of hypothalamic and midbrain galanin. Measurement of tissue NE and MHPG levels in LC projection regions and Fos expression in the LC suggest that the noradrenergic system is largely functioning normally in *Gal^cKO-Dbh^* mice, at least under baseline conditions and following an acute stressor. Finally, we found that *Gal^cKO-Dbh^* mice display a more active coping strategy in three independent tasks measuring active defensive behaviors in an anxiogenic context (marble burying, shock probe defensive burying, novelty suppressed feeding), but their performance in other tests relevant to anxiety, depression, gross motor function, and cognition are indistinguishable from littermate controls.

Prior studies using slice electrophysiology show that galanin has an inhibitory effect on LC neuron activity (Xu et al. 2005; Hökfelt et al. 2018). Additionally, it has been suggested that galanin can be released somatodentritically from large dense core vesicles in LC-NE neurons and thereby produce an autoinhibitory effect on LC neuron activity, although this hypothesis has yet to be tested directly (Huang et al. 2007; Vila-Porcile et al. 2009). While the lack of NE-LC system deficits in the *Gal^cKO-Dbh^* mice may seem surprising, it is important to keep in mind that the measures for noradrenergic system function used in the present study were indirect and may have lacked the sensitivity to detect subtle changes. Although steady-state tissue levels of NE and MHPG have been used to estimate NE turnover, *in vivo* microdialysis is a more direct measure of NE overflow and transmission. Likewise, while Fos is a validated proxy for neuronal activity, it is a binary response that lacks the temporal resolution and sensitivity of electrophysiology. Despite these methodological considerations, our data indicate that the noradrenergic system is not grossly altered in the *Gal^cKO-Dbh^* mice, and that their behavioral phenotypes are caused by galanin depletion rather than alterations in NE signaling. Because *Gal^cKO-Dbh^* mice lack galanin expression in the LC throughout development, it is possible there may be compensatory mechanisms that allow LC activity regulation in the absence of autoinhibitory galanin release. Additionally, our immunohistochemical data uncovered a previously unrecognized galaninergic input to LC neurons that arises from a non-noradrenergic source(s) (**Fig. 4**) that persists in *Gal^cKO-Dbh^* mice and could play a role in regulating LC activity.

Previous studies using the neurotoxin 6-hydroxydopamine (6-OHDA) to lesion the LC suggested that the majority of galanin in the hippocampus and PFC comes from noradrenergic sources by examining the overlap between DBH-positive and galanin-positive fibers or measuring galanin peptide levels in brain regions innervated by the LC (Weiss et al. 1998; Xu et al. 1998; Hökfelt et al. 1998; Melander 1986). However, these experiments had only been performed in rats, and neurotoxin lesioning is not cell type-specific, is rarely complete, can lead to axonal sprouting, and causes neuroinflammation. We overcame these caveats by using region-specific tissue measurements and galaninergic fiber analysis to show that *Gal^cKO-Dbh^* mice have decreased levels of galanin peptide in LC projection fields, including the hippocampal DG and PFC. Our ELISA results indicated that galanin levels in the hippocampus and PFC of the *Gal^cKO-Dbh^* mutants was similar to that observed in mice lacking galanin in all neurons and glia, supporting the idea that galanin in these regions comes primarily from noradrenergic sources (**Fig. 5**). Similarly, fiber analysis showed a significant reduction of galanin-positive fibers in the *Gal^cKO-Dbh^* compared to littermates, although it was still above the level seen in the negative control tissue (**Fig. 6**). This slight divergence may be a result of differences in the sensitivity of these two techniques. Regardless, when taken together these results demonstrate that a significant portion of galanin protein found in the cortex and hippocampal DG comes from noradrenergic sources.

A previous study showed that intracerebroventricular 6-OHDA treatment in rats eliminated DBH-positive fibers in both the dorsal and ventral hippocampus, but abolished galanin-positive fibers in only the dorsal hippocampus, suggesting that galanin in the ventral hippocampus may come from non-noradrenergic neurons (Xu et al. 1998). We observed galanin depletion in both the dorsal and ventral hippocampus of *Gal^cKO-Dbh^* mice, possibly indicative of a species differences in the source of galanin in the ventral hippocampus. Further, our data suggest that, in mice, galanin in the midbrain and hypothalamus is derived from non-noradrenergic sources, consistent with the abundant expression of galanin in many hypothalamic neurons (Skofitsch and Jacobowitz 1985; Melander 1986). By contrast, a previous study in rats reported decreased galanin in the ventral tegmental area following LC lesion using 6-OHDA (Weiss et al. 1998). Our ELISA and fiber analysis results do not support this conclusion, as we saw no change in galanin peptide or galaninergic fibers in the midbrain as a whole or VTA specifically in *Gal^cKO-Dbh^* mice compare to their littermates. This difference may be due to the improved specificity of our knockout model compared to lesion studies, or could indicate a species difference.

The performance of *Gal^cKO-Dbh^* mice was normal in all behavioral assays examined except for ethological-based anxiety tests that measure active, innate patterns of species-typical defensive behaviors. Specifically, *Gal^cKO-Dbh^* mutants buried more marbles, likely a result of increased digging (Thomas et al. 2009), and showed increased digging in the shock probe defensive burying task (**Fig. 9**). Increased digging is sometimes interpreted as increased anxiety-like behavior and could indicate that endogenous noradrenergic-derived galanin may have an anxiolytic effect. However, our other findings do not support this interpretation, as we did not see any differences in canonical approach-avoidance tests for anxiety-like behavior such as the open field, elevated plus maze, and zero maze (**Fig. 8**). These canonical tasks rely on innate, passive avoidance behavior to measure anxiety, whereas marble burying and shock probe burying are defensive tasks that rely on an active response (e.g., digging) as the measure of anxiety behavior, and may elicit increased levels of stress compared to approach-avoidance tasks (Cryan and Sweeney 2011). Increased digging in mice has also been interpreted as a repetitive, perseverative behavior, but we did not observe any other evidence of repetitive behaviors in these mice, such as increased nestlet shredding (Angoa-Perez et al. 2013; Thomas et al. 2009; Angoa-Perez et al. 2012). Additionally, in the novelty-suppressed feeding task, we found that *Gal^cKO-Dbh^* mutants start eating sooner than controls, traditionally interpreted as decreased anxiety- or depressive-like behavior. This task is a conflict task that pits the desire for food against the fear of being in a novel environment, and therefore involves a component of motivation that the other tasks do not possess. Combined, these behavioral data indicate a role for noradrenergic-derived galanin in regulating active coping strategies to environments that elicit immediate or potential threat. To our knowledge, the conventional galanin knockout mice have not been tested in either the canonical or noncanonical anxiety-like behavioral assays used in the present study, but these experiments would allow for an interesting comparison of the role of noradrenergic-derived galanin versus galanin from other sources.

Although the mechanism and circuitry underlying the increase in active coping in the *Gal^cKO-Dbh^* mutants is unclear, both galanin and NE signaling in the amygdala have been implicated in the regulation of stress-related behaviors. For example, intra-amygdala infusion of galanin decreased punished drinking episodes, which could be interpreted as a decrease in active coping (Moller et al. 1999) and is consistent with the enhanced active stress responses we observed in the *Gal^cKO-Dbh^* mutants. Another potential downstream target is the lateral septum, which is innervated by both galaninergic and noradrenergic fibers (Menard and Treit 1996; Melander 1986), and local infusion of the galanin receptor antagonist M40 decreased digging behavior in the shock probe defensive burying test in rats (Echevarria et al. 2005). This result suggests that galanin acting in the lateral septum has the opposite effect of what we report for the *Gal^cKO-Dbh^* mice, but this may be due to a species difference or an alteration in the function of the lateral septum in the *Gal^cKO-Dbh^* mutants due to the lack of galanin throughout development.

Neuropeptides, such as galanin, are preferentially released when neurons fire at high frequencies, and previous research has repeatedly suggested a role for galanin under challenging conditions that strongly activate noradrenergic neurons (Lang et al. 2015; Bartfai et al. 1988; Sciolino and Holmes 2012). A recent study showed that selective optogenetic activation of galanin-containing LC neurons was sufficient to induce avoidance behavior (McCall et al. 2015). β-adrenergic receptor blockade prevented this effect, indicating that LC-derived NE plays a role, but a potential contribution of galanin was not tested. It is possible that stronger behavioral phenotypes would emerge if the noradrenergic systems of *Gal^cKO-Dbh^* mutants were challenged by increasing the intensity and/or duration of stress during or prior to behavioral testing. It may also be informative to test the *Gal^cKO-Dbh^* mutants following chronic voluntary exercise, which is widely reported to increase galanin in the LC and has anxiolytic/antidepressant properties (Sciolino et al. 2012; Sciolino et al. 2015; Epps et al. 2013). Future studies using environmental manipulations, as well as optogenetic strategies for manipulating noradrenergic neuron activity, may help uncover a role for NE-derived galanin in modulating coping behaviors often associated with anxiety and depression.

Collectively, our studies using the *Gal^cKO-Dbh^* mutant model reveal that galanin plays an important role within the NE system by organizing a characteristic repertoire of defensive behaviors. This model is an invaluable tool to probe the many other functions hypothesized to be under the control of NE-derived galanin. Similar strategies targeting other cell type-specific galaninergic populations will help elucidate the broader neuroanatomy of the galanin system and its contribution to many physiological processes and disease states.

## ACKNOWLEDGEMENTS AND AUTHOR CONTRIBUTIONS

We thank Philip V. Holmes, Jessica Hooversmith, Hannah Yoder, and Diane D’Agostin for their technical assistance. Valuable support was provided by the NIEHS Fluorescence Microscopy and Imaging and Transgenic Cores. RPT, NRS, PJ, and DW conceived, designed, and supervised the project. PJ and NWP created the *Dbh^cre^* and *Gal^cKO^* mouse lines. Immunohistochemistry, *in situ* hybridization, and image acquisition was performed by RPT, NRS, KGS, MH, and JMP. Fiber quantification was performed by MH, JMP, and NRS. Behavioral and neurochemical experiments were performed by RPT under the guidance of DW. DL assisted with neurochemical experiments. Mouse husbandry and genotyping at Emory University were performed by CJ. RPT, NRS, PJ, and DW wrote the manuscript with input from co-authors.

